# Translation of *MT-ATP6* pathogenic variants reveals distinct regulatory consequences from the co-translational quality control of mitochondrial protein synthesis

**DOI:** 10.1101/2021.08.30.458165

**Authors:** Kah Ying Ng, Uwe Richter, Christopher B. Jackson, Sara Seneca, Brendan J. Battersby

**Affiliations:** Institute of Biotechnology, University of Helsinki, Helsinki, Finland; Faculty of Biological and Environmental Sciences, University of Helsinki, Helsinki, Finland; Wellcome Centre for Mitochondrial Research, Biosciences Institute, Faculty of Medical Sciences, Newcastle University, Newcastle upon Tyne, UK; Department of Biochemistry and Developmental Biology, Faculty of Medicine, University of Helsinki, Helsinki, Finland; Center for Medical Genetics/Research Center Reproduction and Genetics, Universitair Ziekenhuis Brussel, Vrije Universiteit Brussel (VUB), Brussels, Belgium

## Abstract

Pathogenic variants that disrupt human mitochondrial protein synthesis are associated with a clinically heterogenous group of diseases. Despite an impairment in oxidative phosphorylation being a common phenotype, the underlying molecular pathogenesis is more complex than simply a bioenergetic deficiency. Currently, we have limited mechanistic understanding on the scope by which a primary defect in mitochondrial protein synthesis contributes to organelle dysfunction. Since the proteins encoded in the mitochondrial genome are hydrophobic and need co-translational insertion into a lipid bilayer, responsive quality control mechanisms are required to resolve aberrations that arise with the synthesis of truncated and misfolded proteins. Here, we show that defects in the OXA1L-mediated insertion of MT-ATP6 nascent chains into the mitochondrial inner membrane are rapidly resolved by the AFG3L2 protease complex. Using pathogenic *MT-ATP6* variants, we then reveal discrete steps in this quality control mechanism and the differential functional consequences to mitochondrial gene expression. The inherent ability of a given cell type to recognize and resolve impairments in mitochondrial protein synthesis may in part contribute at the molecular level to the wide clinical spectrum of these disorders.

## Introduction

Human mitochondrial DNA (mtDNA) encodes 13 core protein subunits of the oxidative phosphorylation (OXPHOS) complexes (1). Defects in the faithful synthesis of these proteins represent the largest and most clinically diverse group of human mitochondrial disorders (2). At the molecular level, these disorders are typically characterised by a biochemical defect in OXPHOS, yet the clinical presentation cannot be explained by the bioenergetic dysfunction alone. Understanding the basis by which distinct pathogenic variants impair mitochondrial protein synthesis and organelle homeostasis is critical to determine the molecular pathogenesis of these disorders. The hydrophobicity of these 13 membrane proteins requires co-translational insertion into the inner mitochondrial membrane prior to sorting into distinct assembly pathways for each OXPHOS complex (3,4). Even though a detailed molecular understanding emerges of mitochondrial protein synthesis (5), critical gaps in knowledge nonetheless remain in the quality control mechanisms required to resolve aberrations in protein synthesis (6).

Mitochondrial nascent chain insertion into the inner membrane is mediated by OXA1L (7,8). This protein is a member of the Oxa1 insertase superfamily, which act functionally through a conserved mechanism whereby the insertase forms a positively charged groove to facilitate an interaction via the short hydrophilic sequence of substrates which precede the transmembrane domain (9,10). As a consequence, this facilitates the translocation of substrates across the bilayer to obtain their correct topology. Biallelic pathogenic variants in human *OXA1L* present with severe encephalopathy, hypotonia and developmental delay and a defect in OXPHOS complex assembly (11). A similar molecular phenotype is also observed with reverse genetic approaches to silence OXA1L function in cultured human cells (12). In yeast, OXA1 also mediates the N-terminal insertion of nuclear-encoded substrates into the inner membrane of the organelle (13). In terms of mitochondrially-encoded substrates, the most extensive investigations have been conducted in the budding yeast with Cox2 of cytochrome c oxidase and Atp6 and Atp9 of the F_1_F_O_ ATP synthase (7,8,14). Biochemical data has long supported contact between the C-terminal extension of OXA1L and mitochondrial ribosomes (15,16), suggesting a co-translational interaction. Recently, a high-resolution structure identified the molecular basis of this mechanism. OXA1L binds the mitochondrial ribosome at three sites that correlates with conformational changes in mL45 to limit folding of the emerging nascent chain in the exit tunnel (17). Despite the co-translational contact between OXA1L and the mitochondrial ribosome, a gap remains between the exit site and the membrane that could allow additional interactions with the emerging nascent chains prior to membrane insertion.

Following insertion, newly synthesized polypeptides follow two fates: stable assembly into an OXPHOS complex or degradation. The AAA (<u>A</u>TPases <u>a</u>ssociated with various cellular <u>a</u>ctivities) protease complex composed of AFG3L2 subunits plays a key role in the quality control of mitochondrial nascent chains (18–21). In humans, AFG3L2 exists as a homo- or hetero-hexameric complex with paraplegin subunits anchored in the inner membrane and facing the matrix space. Pathogenic variants in these subunits manifest as autosomal dominant and or recessive neuromuscular disorders (22–27) and appear to cluster into distinct structural elements of the complex (28). Progressive loss of AFG3L2 function triggers a stress response that remodels mitochondrial membrane morphology and a series of pathways that negatively affect mitochondrial gene expression (20,21,29). In the budding yeast there is a genetic interaction between this mitochondrial matrix AAA protease complex, the OXA1 insertase and assembly of the F_1_F_O_ ATP synthase (30), where the chaperone function of the complex appears to be particularly important (31). Despite a detailed understanding on the structural mechanisms by which the AFG3L2 complex mediates protein unfolding and proteolytic degradation (28), the basis and selectivity by which mitochondrial nascent chain substrates are delivered to the complex remains poorly understood.

Currently, little is known about the regulation of co-translation quality control mechanisms required to resolve aberrations in mitochondrial protein synthesis (6). Although there is spatial separation of the mitochondrial genome from RNA processing (32), there is no physical barrier within the organelle to enforce strict quality control mechanisms to prevent translation of faulty mRNA, such as truncated transcripts. As a consequence, the importance of co-translational quality control mechanisms is paramount for organelle homeostasis and cell fitness.

In this study, we establish a model by which failures in the co-translational insertion of MT-ATP6 are rapidly resolved by the AFG3L2 complex. Using specific *MT-ATP6* pathogenic variants, we then reveal discrete regulatory steps in the quality control of mitochondrial nascent chain synthesis and the phenotypic consequences from disruptions in these processes. Collectively, our findings point to the differential effects that translation of mitochondrial pathogenic variants of the same gene can exert on organelle homeostasis.

## Results

### MT-ATP6 as a model substrate for the study of co-translational quality control

Despite the importance of OXA1L and AFG3L2 to OXPHOS complex assembly, how these two factors work synergistically during mitochondrial protein synthesis remains unclear. To investigate these functions, we performed single and combined siRNA knockdown of *AFG3L2* and *OXA1L* in human cultured fibroblasts followed by metabolic labelling with ^35^S methionine/cysteine (Fig. 1A-C). A robust inhibition of OXA1L function did not impair the overall synthesis of mitochondrial nascent chains (Fig. 1B). However, the MT-ATP6 and MT-CO2 nascent chains were completely unstable and rapidly degraded, whereas the stability of other mitochondrial nascent chains was not adversely affected from OXA1L inhibition in this assay (Fig. 1B, C). We have shown previously that acute pharmacological inhibition of AFG3L2 can lead to the accumulation of mitochondrial nascent chains (20). Therefore, we addressed whether AFG3L2 was responsible for the rapid degradation of MT-ATP6 and MT-CO2 nascent chains. The double knockdown of *AFG3L2* and *OXA1L* did not impair nascent chain synthesis but did prevent the rapid degradation of both MT-ATP6 and MT-CO2 (Fig. 1B, C). Although these two mitochondrially-encoded proteins differ considerably in their structure and hydrophobicity (33,34), the apparent inability to insert these nascent chains into the inner membrane induces proteolytic degradation by the AFG3L2 complex. Together, the data establish MT-ATP6 as model substrate with which to investigate co-translation quality control mechanisms of mitochondrial protein synthesis.

**Figure 1.**
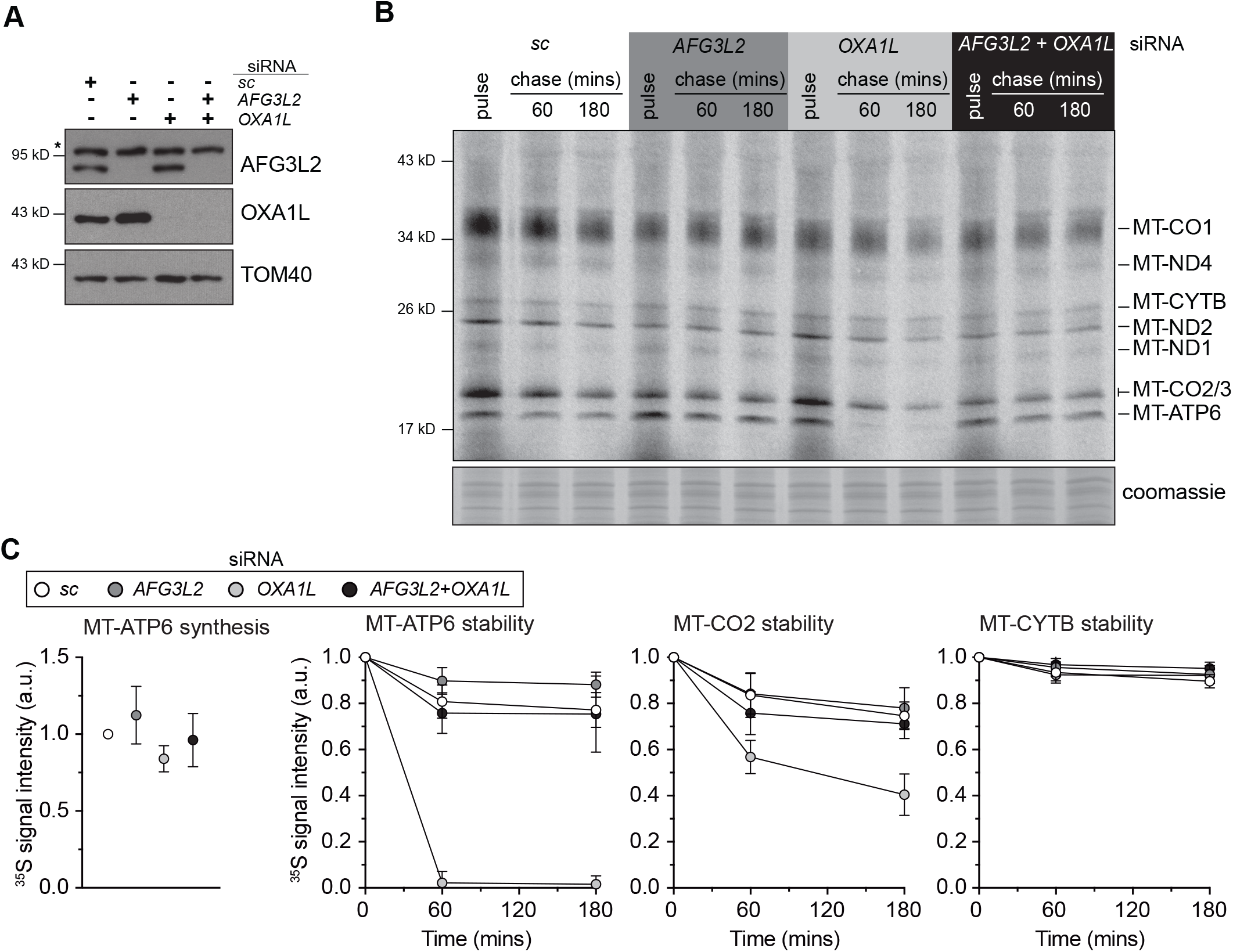
MT-ATP6 as a model substrate for the study of co-translational quality control. (**A**) Immunoblotting of whole-cell lysates from human fibroblasts treated with the indicated siRNAs. sc, scrambled control. *Nonspecific band detected with AFG3L2 antibody. (**B**) A representative image of ^35^S-methionine/cysteine metabolic labelling of mitochondrial protein synthesis in human fibroblasts treated with the indicated siRNAs. Cells were pulse-labelled for 30 minutes, followed by 60 min and 180 min cold chase. (**C**) Quantification of the metabolic labelling from five independent experiments with the mean ± SD.

### Pathogenic variants of MT-ATP6 to probe mitochondrial co-translation quality control

To investigate the co-translation quality control of MT-ATP6, we turned to two pathogenic mtDNA variants segregating in patient-derived skin fibroblasts that distinctly affect the translation of this open reading frame (ORF) (Fig. 2A). The first variant (m.9205delTA) encodes a 2-bp deletion that removes the in-frame stop codon required for translation termination of MT-ATP6 (35) and will be referred to as the non-stop mRNA. The second variant (m.8611insC, L29PfsX36) encodes a 1-bp insertion that alters the *MT-ATP6* reading frame, leading to premature translation termination after 36 amino acids (36) and will be referred to as the frameshift (FS) mRNA. Both variants are present at high heteroplasmy levels (e.g., 97% and 80%, respectively) in cultured fibroblasts (21).

**Figure 2.**
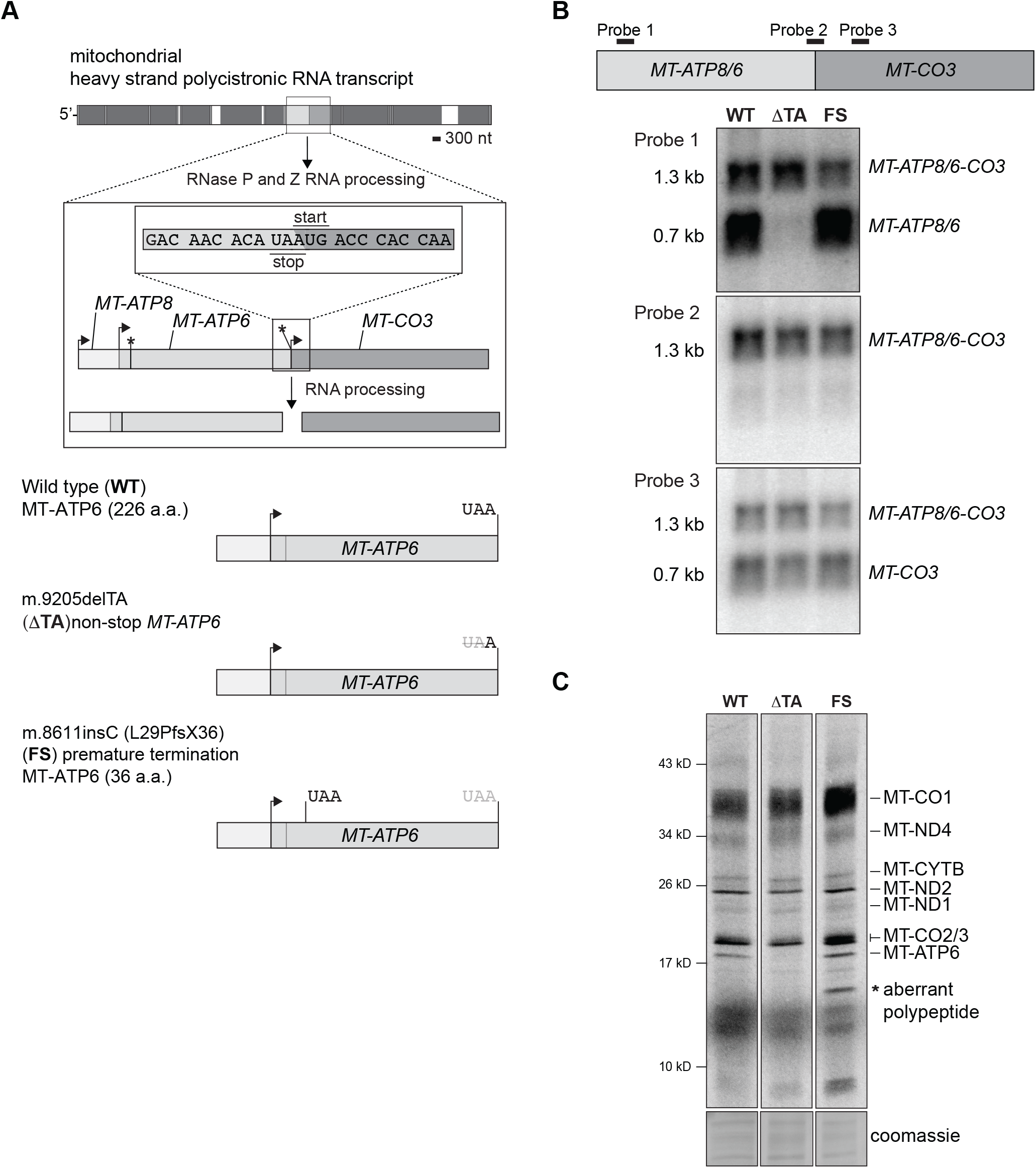
The effect of pathogenic *MT-ATP6* variants at the RNA level and on mitochondrial protein synthesis. (**A**) Top, a schematic illustrating the RNA processing of the mitochondrial polycistronic transcript to liberate MT-ATP6 mRNA. Arrows indicate the position of the AUG start codon and asterisk the respective stop codon. *MT-ATP8* and *MT-ATP6* have an overlapping start and stop codons. *MT-CO3* does not encode a stop codon. (**B**) Top, diagram illustrating the binding sites of oligonucleotide probes used in northern blotting analysis. Northern blotting of total whole cell RNA isolated from cultured human fibroblasts with the indicated *MT-ATP6* genotypes. Each panel represent a representative image following hybridization with the indicated oligonucleotide probes. (**C**) A representative 30-minute pulse of metabolic labelling with ^35^S-methionine/cysteine of the cultured human fibroblasts with the indicated *MT-ATP6* genotypes. All data are representative of independent experiments.

Within human mtDNA, *MT-ATP6* has an overlapping open reading frame with *MT-ATP8* (1). At the mRNA level, two transcripts can be detected for *MT-ATP6*. The first is a tricistronic mRNA that contains *MT-ATP8, MT-ATP6* and *MT-CO3* and a shorter processed transcript of *MT-ATP8* and *MT-ATP6* (Fig. 2A, B). In human wild type fibroblasts, both mRNAs are abundantly detected (Fig. 2B). The RNA processing mechanism of the tricistronic transcript has not yet been established but it is not due to the canonical endonucleolytic cleavage catalysed by the mitochondrial RNase P complex and RNase Z (37). Surprisingly in the m.9205delTA patient fibroblasts, we could not detect the shorter *MT-ATP6* mRNA transcript with northern blotting of total RNA nor MT-ATP6 synthesis with metabolic labelling (Fig. 2B, C). In contrast, the m.8611insC patient fibroblasts had no difference in the abundance of the two *MT-ATP6* mRNA transcripts or synthesis of the corresponding nascent chain (Fig. 2B, C). Although, an aberrant mitochondrial polypeptide was generated in the pulse metabolic labelling of fibroblasts segregating the *MT-ATP6* frameshift variant (Fig. 2C). Despite the fact that two *MT-ATP6* transcripts are readily detected in human cultured cells, it has not been established which transcript is used as a template in mitochondrial protein synthesis.

### The tricistronic transcript is associated with mitochondrial ribosomes

To address this question, we sought to determine which *MT-ATP6* mRNA is associated with mitochondrial ribosomes using the wild type and m.9205delTA fibroblasts. Mitochondrial ribosomes were isolated by sucrose density gradient preparation that revealed no difference in the assembly profiles (Fig. 3A). Northern blotting of pooled fractions isolated from this preparation demonstrate that the longer tricistronic mRNA containing *MT-ATP8/6/CO3* is the predominant transcript associated with the 55S mitochondrial monosome in both wild type and m.9205delTA fibroblasts (Fig. 3B). In contrast, the shorter *MT-ATP6* and *MT-CO3* transcripts clearly appear to sediment in lighter fractions of the sucrose gradient.

**Figure 3.**
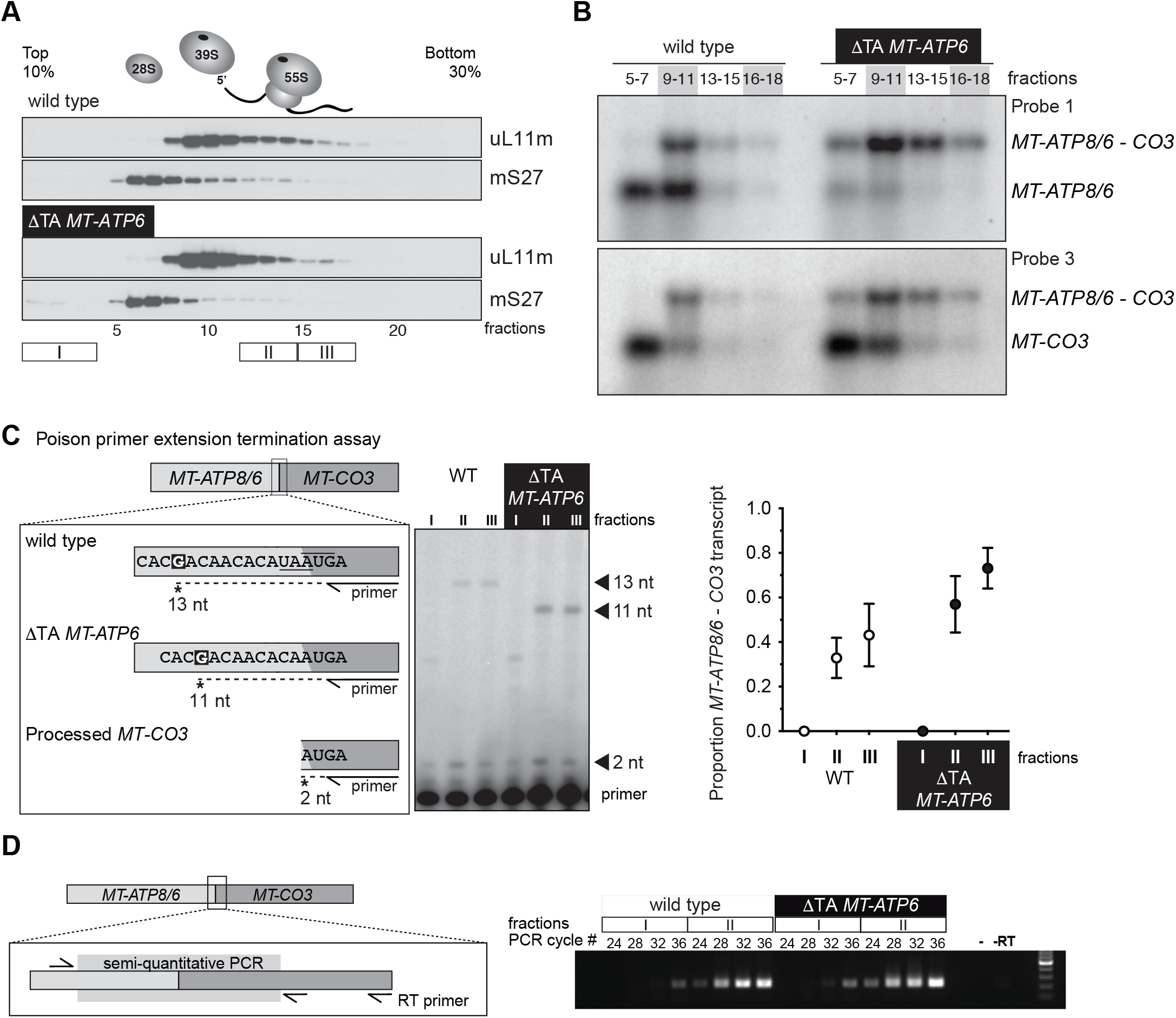
The longer tricistronic transcript of *MT-ATP8, MT-ATP6 and MT-CO3* associates with the mitochondrial ribosome. (**A**) Immunoblotting of sucrose density gradient fractions of the cultured human fibroblasts with the indicated *MT-ATP6* genotypes to profile the sedimentation profile of the mitochondrial ribosome. (**B**) Northern blotting of RNA isolated from the indicated sucrose density gradient fractions. (**C**) Left, a diagram of the poison primer extension assay to determine the tricistronic transcript from processed *MT-CO3* in the indicated *MT-ATP6* genotypes. Middle, a representative poison primer extension analysis from total RNA collected from the indicated sucrose gradient fractions. Right, quantification of the primer extension analysis from three independent experiments (mean +/- S.D.) with the indicated *MT-ATP6* genotypes. (**D**) Semi-quantitative RT-PCR of RNA isolated from the indicated sucrose gradient fractions of cultured human fibroblasts with the indicated *MT-ATP6* genotypes. Left, a diagram of the assay. Right, representative image of the analysis. -, PCR with no cDNA template; -RT, PCR without the reverse transcriptase step. All data are representative of independent experiments.

Next, we turned to two orthogonal approaches to further validate our northern blotting results. In the first, we developed a poisoned primer extension assay to provide single nucleotide resolution of the *MT-ATP6* mRNAs. A primer annealing to *MT-CO3* was radiolabelled at the 5’ end to be used for reverse transcription. In this assay, cDNA synthesis terminates following incorporation of the chain-terminating dideoxynucleotide ddCTP when a guanine nucleotide is encountered in the template sequence (Fig. 3C). The assay distinguishes the 2-bp deletion in the *MT-ATP6* mRNA associated with the m.9205delTA fibroblasts (Fig. 3C). In both wild type and the m.9205delTA fibroblasts, the longer tricistronic transcript associates with mitochondrial ribosomes whereas the shorter processed *MT-CO3* transcript is enriched at the top of the gradient (Fig. 3C). Lastly, we used strand-specific reverse transcription followed by semi-quantitative PCR, which also showed an enrichment of the *MT-ATP8/6/CO3* transcript in the monosome fractions of both wild-type and m.9205delTA fibroblasts (Fig. 3D).

Although the m.9205delTA *MT-ATP6* variant was originally described as a non-stop mRNA (38), the tricistronic transcript with a 2-bp deletion generates an in-frame fusion ORF with *MT-ATP6* and *MT-CO3*. To test such a hypothesis, we developed an *in vitro* translation assay. Since a completely reconstituted translation system does not yet exist for human mitochondria, we used the bacterial PURE system. We took the sequence flanking the junction of wild type human *MT-ATP6* and *MT-CO3* to insert it between two bacterial genes, *DHFR* and *bla* (Fig. 4A). The stop codon in the upstream *DHFR* was removed so that translation termination would be dependent upon the UAA within the *MT-ATP6* sequence. An upstream ribosomal binding site was placed 5’ to the DHFR ORF, thus translation will be skewed towards the synthesis of this product. We also generated two variations of the construct, one contained an additional adenine (+A) to produce distinct stop and start codons while the other contained the 2-bp deletion found in m.9205delTA (ΔTA) (Fig. 4A). As expected, in this *in vitro* translation assay the stop codon in the wild type and +A constructs favours synthesis of the upstream DHFR whereas the ΔTA template generates a fusion ORF (Fig. 4B). Together, the data provide robust evidence that the absence of a stop codon in the m.9205delTA variant would generate a fusion ORF between *MT-ATP6* and *MT-CO3*.

**Figure 4.**
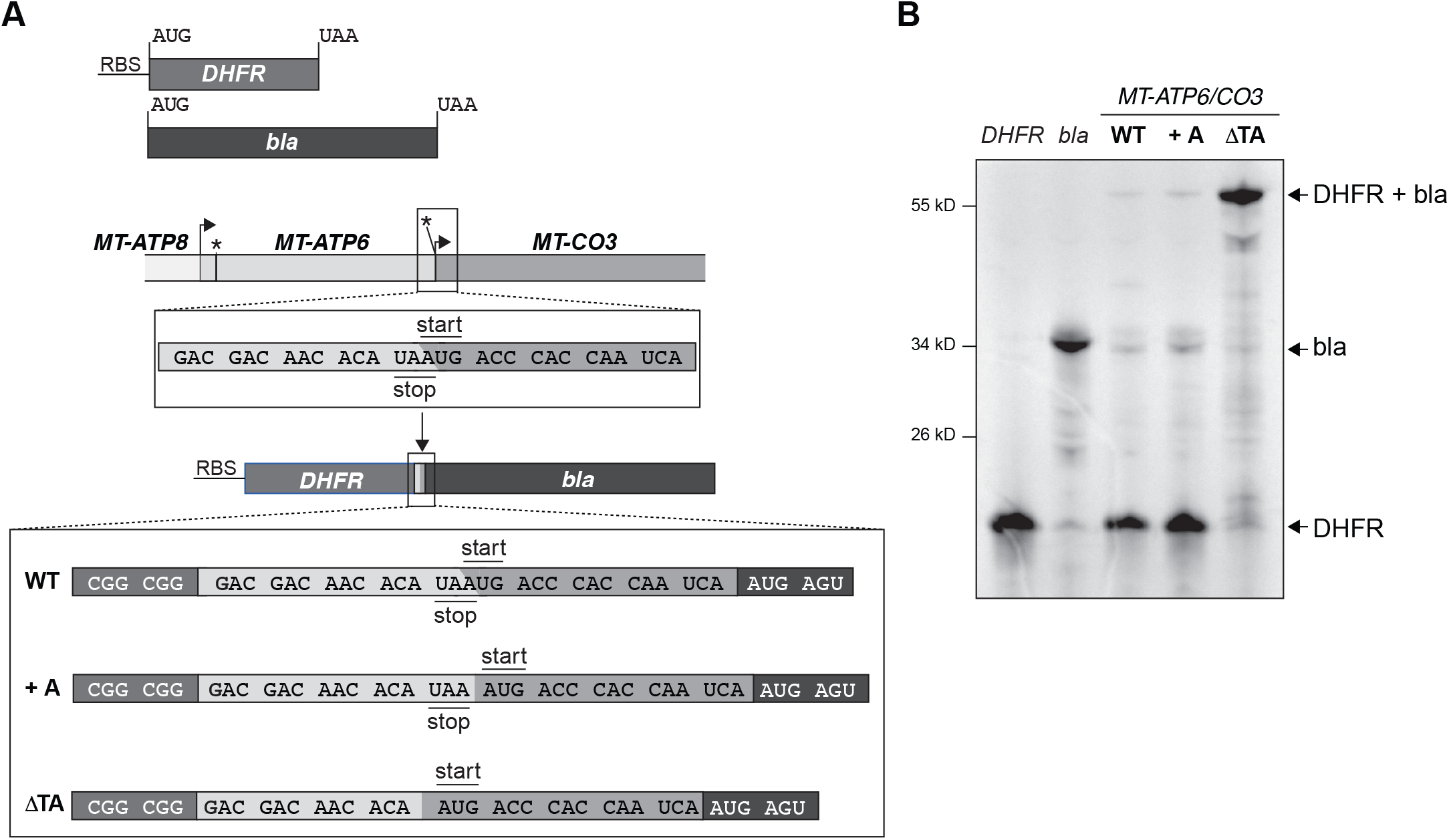
An *in vitro* translation assay to evaluate the synthesis of *MT-ATP6* variants. (**A**) A schematic illustrating the template design. +A, an additional adenine nucleotide was added to generate a stop codon. ΔTA, represent the m.9205delTA *MT-ATP6* variant. (**B**) Autoradiogram of a representative *in vitro* experiment using the indicated templates with ^35^S-methionine/cysteine to label all nascent chains following a 90-minute incubation. All samples were treated with a nuclease prior to gel loading to hydrolyse any potential polypeptidyl tRNAs.

### Differential effects on mitochondrial protein synthesis from the loss of co-translational quality control of MT-ATP6 pathogenic variants

The two pathogenic *MT-ATP6* variants provide substrates to test mechanisms in co-translational quality control of mitochondrial protein synthesis using the model established in Fig 1. The fusion ORF encoded by the m.9205delTA variant will generate an aberrant polypeptide with impaired membrane insertion and will misfold during synthesis. In contrast, the frameshift mutation will generate a short polypeptide of 36 amino acids that will be too short to engage co-translationally with the OXA1L insertase. First, we used the m.9205delTA variant with siRNA knockdown of *AFG3L2* and *OXA1L* to assess *de novo* mitochondrial protein synthesis. Unlike in wild type fibroblasts, to our surprise, loss of either the OXA1L insertase or the AFG3L2 complex induced a profound inhibitory effect on the synthesis of all mitochondrial nascent chains (Fig. 5B, C). This suggests that failures in co-translational protein membrane insertion and or degradation following translation of the MT-ATP6 m.9205delTA variant generates negative feedback onto mitochondrial protein synthesis. Next, we turned to the m.8611insC variant focusing on the role of AFG3L2. In this background, loss of function for the AFG3L2 complex induced a similar phenotype to what we observed in wild type fibroblasts (Fig. 1,6). Further, the abundance of the aberrant polypeptide generated in the m.8611insC fibroblasts (Fig. 2C) was unaffected with the loss of AFG3L2. Despite both *MT-ATP6* pathogenic variants manifesting with a biochemical defect in OXPHOS (21,39), our data point to distinct molecular consequences from the translation of these pathogenic mRNAs and the quality control requirements in mitochondrial gene expression.

**Figure 5.**
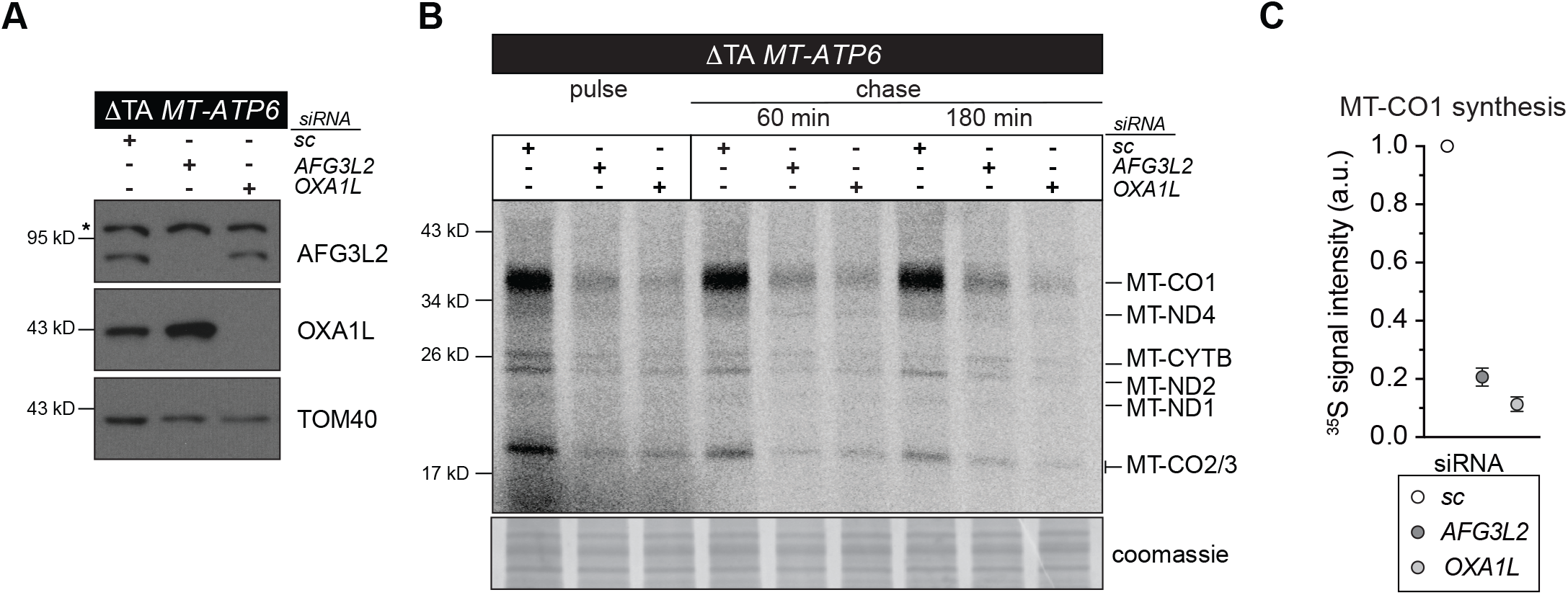
Defective co-translational insertion and quality control of the *MT-*ATP6 fusion open reading frame encoded by the m.9205delTA variant impairs mitochondrial protein synthesis. (**A**) Immunoblotting of whole-cell lysates from cultured human fibroblasts segregating the m.9205delTA variant following treatment with the indicated siRNAs. sc, scrambled control; asterisk, nonspecific band detected with AFG3L2 antibody. (**B**) A representative image of ^35^S-methionine/cysteine metabolic labelling of mitochondrial protein synthesis in human fibroblasts segregating the m.9205delTA variant treated with the indicated siRNAs. Cells were pulse-labelled for 30 minutes, followed by 60 min and 180 min cold chase. (**C**) Quantification of the metabolic labelling from three independent experiments with the mean ± SD.

**Figure 6.**
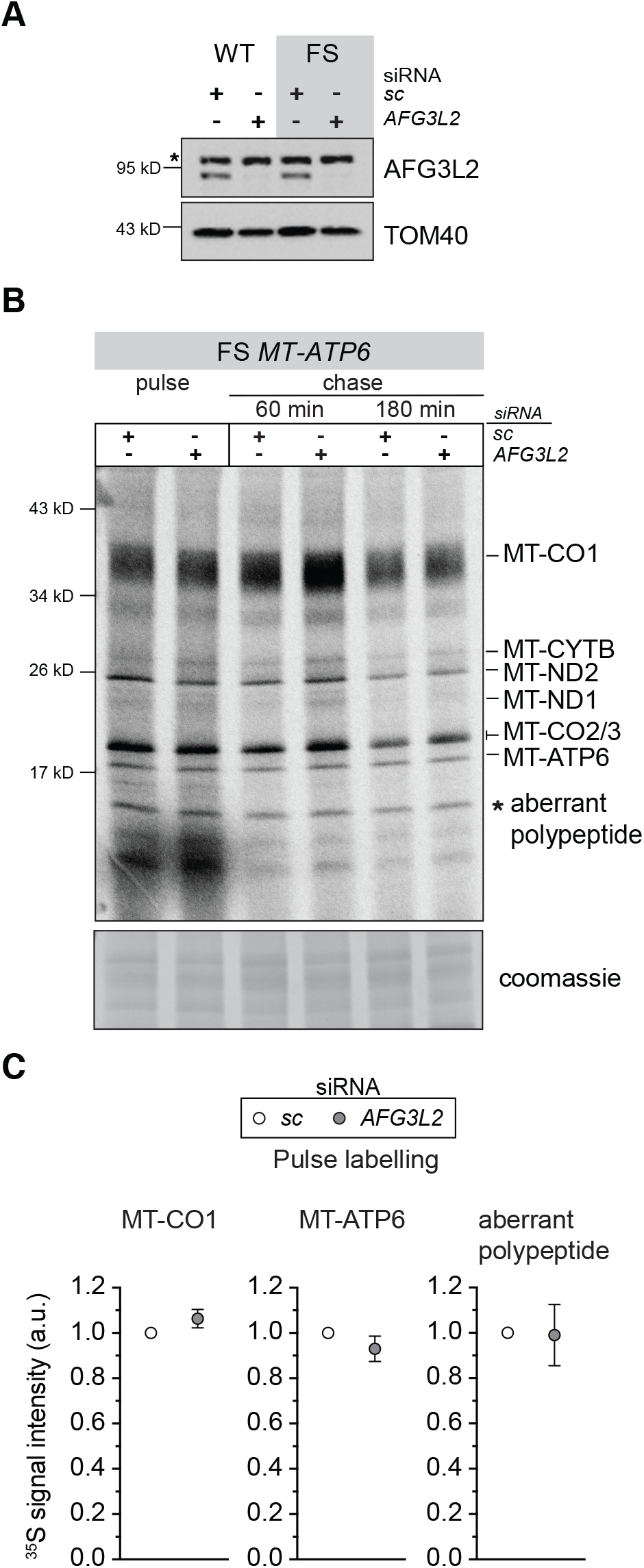
Loss of AFG3L2 function with translation of the m.8611insC *MT-ATP6* frameshift variant does not adversely affect mitochondrial protein synthesis. (**A**) Immunoblotting of whole-cell lysates from cultured human fibroblasts with the indicated *MT-ATP6* genotypes following treatment with the indicated siRNAs. sc, scrambled control; asterisk, nonspecific band detected with AFG3L2 antibody. (**B**) A representative image of ^35^S-methionine/cysteine metabolic labelling of mitochondrial protein synthesis in human fibroblasts segregating the m.8611insC variant treated with the indicated siRNAs. Cells were pulse-labelled for 30 minutes, followed by 60 min and 180 min cold chase. (**C**) Quantification of the metabolic labelling from three independent experiments with the mean ± SD.

## Discussion

Disruptions to protein synthesis are one of the most common causes of human mitochondrial diseases but yet present with tremendous clinical heterogeneity (2). For a long-time, these pathogenic variants have been investigated through a prism focusing specifically on the bioenergetic defect arising from the failure in OXPHOS complex assembly. Unfortunately, this perspective overlooks the myriad of ways in which a primary molecular defect in protein synthesis can impinge upon organelle function. Here, we reveal how disruptions to a co-translational step in nascent chain synthesis and quality control can exert differential functional consequences from translation of pathogenic variants in the same gene.

In the fully assembled F_1_F_O_ ATP synthase, the MT-ATP6 subunit has five alpha helices embedded within the membrane, including one formed at the N-terminus (33). Four helices create the critical functional interface with the c-ring, forming the water half-channels for proton translocation. In the absence of co-translational insertion into the membrane, the hydrophobicity of MT-ATP6 nascent chains would likely generate protein misfolding upon exit of the mitochondrial ribosome. This nascent chain misfolded state is clearly recognized to initiate proteolytic degradation but how it is facilitated is not so clearly understood. The AFG3L2 complex requires substrates to possess unstructured N or C-termini at least 20 amino acids in length (40). Our previous research using the peptidomimetic actinonin, which resembles formylated methionine, established the small molecule as a robust inhibitor of the AFG3L2 complex (20). Since mitochondrial translation initiation requires a formyl methionine (41), a mechanism consistent with the data would suggest that failures in the membrane insertion of MT-ATP6 will be delivered to the AFG3L2 complex via the N-terminus. Considering the spatial constraints between the ribosome and membrane (17), however, it is likely that additional factors first recognize the misfolded protein to facilitate subsequent delivery to the AFG3L2 complex. The use of proximity labelling approaches to identify spatial functional domains or networks within mitochondria (42,43) will likely be critical to elucidate these mechanisms.

Translation of the MT-ATP6-CO3 fusion ORF will generate a misfolded protein upon the exit of the ribosome. This aberrant nascent chain will disrupt the insertion process and translocation into the membrane. In the metabolic labelling experiments, we did not observe the synthesis of any aberrantly sized protein or polypeptide fragments that could be attributable to this fusion ORF. This could imply rapid degradation. Equally possible, however, is that the synthesized product co-migrates with MT-CO1 as it is predicted to be a similar size and would thus be difficult to distinguish in this assay. The profound inhibitory effect observed in the patient fibroblasts in metabolic labelling when OXA1L and AFG3L2 are missing, suggest that the co-translational quality control of this aberrant protein is paramount for organelle homeostasis. Previously, we established that translation of the m.9205delTA pathogenic variant upon an acute heat shock generated a rapid attenuation in mitochondrial protein synthesis, triggering a ribosome and RNA quality control response but with no effect on mtDNA abundance (21). At the time, we considered the variant as a non-stop mRNA, but our data here clearly demonstrate it is a fusion ORF associated with mitochondrial ribosomes. Since temperature is a well-known modulator for protein folding (44), the findings suggest a mechanism by which a severe proteotoxicity can be induced by translation of a fusion ORF. Our findings may then have relevance to other pathogenic mtDNA variants, such as genome deletions that generate fusion ORFs (45), and the role of acute febrile infections that are well-known to modulate the natural history of these mitochondrial disorders (46). Therefore, understanding the molecular basis by which defects in mitochondrial protein synthesis arise and are resolved across different cell types appears to be a critical modulator to the pathogenesis.

Translation of the *MT-ATP6* frameshift variant will generate a nascent chain 36 amino acids in length. Since the length of the exit tunnel in the human mitochondrial ribosome from the peptidyl transferase centre is approximately 100 Å and prevents secondary helix formation (17), this would correspond to approximately 30 amino acids of a linear polypeptide chain, assuming 3.35 Å per amino acid. Although, there is a gap between the ribosome and OXA1L in the membrane, it is likely that this nascent chain prematurely terminates synthesis prior to becoming fully engaged as a substrate for insertion. In contrast to the data with m.9205delTA pathogenic variant, loss of AFG3L2 function with the frameshift variant exhibited no negative inhibitory feedback onto mitochondrial protein synthesis. This reveals differential consequences from translation of aberrant proteins and the necessary quality control steps to resolve these errors.

Another surprising discovery in our study was the demonstration that the longer tricistronic mRNA encoding *MT-ATP8/6/CO3* is the predominant transcript associated with the 55S mitochondrial monosome in both wild type and patient fibroblasts. The processed shorter transcripts appear enriched off the ribosome. A previous study hinted at such an interpretation. Genetic silencing of FASTKD5 impaired the RNA processing of the tricistronic mRNA but had no effect on MT-ATP6 synthesis or assembly of the F_1_F_O_ ATP synthase (47). Although FASTKD5 binds mitochondrial RNA it is not known to possess any nucleolytic activity, suggesting that other factors are required for the RNA processing event and that this may be translationally regulated. Future studies should reveal the basis of this mechanism.

In summary, our findings highlight the importance of co-translational quality control of mitochondrial protein synthesis and the differential consequences on organelle homeostasis. Going forward, it is important to understand the regulation of these mechanisms across the affected cell types in patients to determine the scope by which primary molecular defects in mitochondrial protein synthesis impair cell fitness.

## Materials and Methods

### Cell culture

Human and mouse fibroblasts were cultured at 37°C and 5% CO_2_ in Dulbecco’s modified Eagle’s medium (Lonza) with high glucose supplemented with 10% foetal bovine serum, 1x glutamax and 50 μg/ml uridine. All primary cells were immortalized by expressing E7 and hTERT (47) unless previously immortalized by the donating laboratory. All cells tested negative for mycoplasma infection (PromoKine). Stealth siRNA against *AFG3L2* (HSS116886), *OXA1L* (HSS107511, MSS229504) and scrambled sequences were obtained from Life Technologies. siRNAs were transfected on day 1 and 3 with LipofectamineTM RNAiMAX (Thermo Fisher) and collected on day 8 for analysis. All siRNA knockdowns were evaluated by immunoblotting using specific antibodies.

### Immunoblotting

Cells were solubilized in phosphate buffered saline, 1% dodecyl-maltoside (DDM), 1 mM PMSF (phenylmethylsulfonyl fluoride), and complete protease inhibitor (Thermo Fisher). Protein concentrations were measured by the Bradford assay (BioRad). Equal amounts of proteins were separated by Tris-Glycine SDS-PAGE and transferred to nitrocellulose membrane by semi-dry transfer. Membranes were blocked in TBST with 1% milk at room temperature for 1 h, followed by incubation with primary antibodies overnight at 4°C in 5% BSA/TBST. Signals were detected the following day with secondary HRP conjugates (Jackson ImmunoResearch) using ECL with x-ray film and iBright Imaging System (Thermo Fisher). Primary antibodies from Proteintech Group: AFG3L2 (14631-1-AP, 1:5000); uL11m (15543-1-AP, 1:20000); mS27 (17280-1-AP, 1:5000); OXA1L (21055-1-AP, 1:5000). Primary antibodies from Abcam/Mitosciences: MT-CO1 (1D6E1A8, 1:500). Primary antibody from Santa Cruz: TOM40 (sc-11414, 1:5000). Representative data of independent experiments were cropped in Adobe Photoshop with only linear corrections to brightness applied.

### Metabolic labelling of mitochondrial protein synthesis

Mitochondrial protein synthesis was analysed by metabolic labelling with ^35^S methionine/cysteine (20). Cells were pre-treated with anisomycin (100 ug/ml) for 5 min to inhibit cytoplasmic translation then pulsed with 200 uCi/ml 35S Met-Cys (EasyTag-Perkin Elmer). In chase experiments, cells were pulsed for 30 min with radioisotope and medium was replaced with one without the radioisotope for the indicated time. Equal amounts of sample protein were first treated with Benzonase (Thermo Fisher) on ice for 20 min and resuspended in 1x translation loading buffer (186 mM Tris-Cl pH6.7, 15% glycerol, 2% SDS, 0.5 mg/ml bromophenol blue, 6% beta-mercaptoethanol). A 12-20% gradient Tris-Glycine SDS-PAGE was used to separate labelled proteins and dried for exposure with a phosphorimager screen and scanned with a Typhoon 9400 or Typhoon FLA 7000 (GE Healthcare) for quantification. Gels were rehydrated in water and stained Coomassie-blue to verify sample loading.

### Isokinetic sucrose gradients

Cells were cultured on 150 mm plates then rapidly transferred to ice where the media was removed, and cells washed with cold PBS. Cells were lysed (50 mM Tris, pH 7.2, 10 mM Mg (Ac)_2_, 40 mM NH_4_Cl 100 mM KCl, 1% DDM, 1 mM ATP, 400 μg/ml chloramphenicol, and 1 mM PMSF) and incubated on ice for 20 minutes. Cell lysates were clarified following centrifugation for 10 minutes at 20 000 x g at 4°C then protein concentrations measured (Bradford). From each cell lysate, a total of 1 mg of protein was loaded on top of a 16 ml linear 10-30% sucrose gradient (50 mM Tris, pH 7.2, 10 mM Mg (Ac)_2_, 40 mM NH_4_Cl 100 mM KCl, 1 mM ATP and 1 mM PMSF) and centrifuged for 15 h at 4°C and 74 400 x g (Beckman SW 32.1 Ti). From the gradient, 24 equal volume fractions were collected for either protein or RNA isolation. Samples for protein analysis were precipitated with TCA. For RNA isolation, fractions were combined according to the established sedimentation profile of mitochondrial ribosomes then concentrated using Microsep™ centrifugal filter (MWCO 10kD) at 7500 x g for 90 min at 4°C. RNA was isolated using TRIzol LS reagent (Thermo Fisher).

### RNA isolation and northern blotting

Total RNA from cultured cells and sucrose gradient fractions were isolated with TRIzol according to the manufacturer’s instructions. All RNA samples to be used in reverse transcriptase (RT) reactions were treated with turbo DNase (Thermo Fisher) followed by re-isolation with TRIzol. All samples were re-precipitated with 0.1 volume of 3M sodium acetate and 3 volume of ice cold 100% ethanol. For Northern blotting, 5 μg of total RNA from each sample was analysed through 1.2% agarose-formaldehyde gels and transferred to HybondTM-N+ membrane (GE Healthcare) by neutral transfer. T4 Polynucleotide Kinase (NEB) 5’ radiolabelled oligonucleotides were used for detection of mitochondrial transcripts (MT-ATP8/6 5’-TGGGTGATGAGGAATAGTGTAAGGAG; MT-CO3 5’-ATAGGCATGTGATTGGTGGGTCAT; MT-ATP6 - MT-CO3 border 5’-ATTGGTGGGTCATTATGTGTTGTC). Hybridization (25% Formamide, 7% SDS, 1% BSA, 0.25M sodium phosphate pH 7.2, 1mM EDTA pH 8.0, 0.25M NaCl2) was performed overnight at 37°C. Membranes were washed (2x SSC/0.1% SDS) and then dried for exposure on a phosphorimager screen (GE Healthcare) and scanned with a Typhoon 9400 (GE Healthcare).

### Primer extension assay

An RNA oligonucleotide (5’-GGCATGTGATTGGTGGGTC) was 5’ labelled with gamma ^32^P ATP by T4 Polynucleotide Kinase (NEB) according to the manufacturer’s instructions. Labelled oligonucleotides were separated from non-incorporated nucleotides via G50 MicroSpin columns (GE Healthcare). 1 μg of total RNA from the indicated gradient fractions were annealed to 0.3 pmol of radiolabelled specific primer in 30 mM Tris-Cl pH 7.5 and 2 mM KCl at 90°C for 2 min and then cooled down to room temperature for 5 min. Reverse transcription reaction with 0.2 U/μL AMV Reverse Transcriptase (NEB) was performed at 42°C for 30 min using a dNTP/ddCTP mix at 0.25 mM each (final concentration). Subsequently, RNaseH (Epicentre) was added to the reaction and incubated at 37°C for 10 min to degrade RNA. To 5 μL cDNA, 5 μL RNA Loading dye (Formamide + bromophenol blue) was added, and RT products were resolved on a 12% denaturing PAGE (7 M Urea,1x TBE, 30cm). After the run, the gel was fixed and dried on Whatman paper and exposed to a phosphorimager screen.

### Semi-quantitative PCR

Maxima H Minus RT (Thermo Scientific) was used according to manufacturer’s instructions. Briefly, the gene specific primer (5’-TGTGCTTTCTCGTGTTACATCGC) was annealed to 300 ng of RNA and reverse transcribed at 50°C for 30 min. For the subsequent PCR the Econo-Taq (Lucigen) master mix was used with forward (5’-TGACTATCCTAGAAATCGCTGTCG) and reverse (5’-GTATGAGGAGCGTTATGGAGTGG) primers that amplify a fragment of *MT-ATP6* and *MT-CO3* border.

### *In vitro* translation

The PURExpress ΔRF123 cell-free translation system (NEB) was used with the following modifications. DNA templates were generated by PCR using the supplied dihydrofolate reductase (DHFR) plasmid as a template, including the *bla* gene. Each construct possessed an upstream T7 promoter sequence and a Shine-Delgarno sequence of the DHFR ORF. In a 20 μl reaction, 1 μl of 100 ng DHFR plasmid was amplified with 0.4 units of Phusion™ High-Fidelity DNA polymerase (Thermo Fisher), 4 μl of 5x Phusion™ HF buffer, 0.4 μl of 10 mM dNTPs, 1 μl of 10 µM primer (forward and reverse) at 95 °C for 5 min, followed by 30 cycles of 95 °C 30 sec, 55 °C 30 sec, 68 °C 30 sec, final extension at 68 °C 7 min and held at 4 °C. PCR product (3 µl) was analysed on 1.5 % agarose-TAE gel and remaining product was purified with NucleoSpin Gel and PCR Clean-up kit (Macherey Nagel). All PCR-generated DNA templates were verified by Sanger sequencing. Equimolar reactions of each specific construct were prepared for *in vitro* transcription and translation reactions in a total volume of 12.5 μl containing 5 μl of PURExpress kit solution A, 3.75 μl of solution B, 0.25 μl each of the supplied release factor (RF1, RF2, and RF3), 8 U of RiboLock RNase inhibitor (Thermo Fisher), 3 ng DNA template, 5 μM anti-ssrA oligonucleotide (5’-TTAAGCTGCTAAAGCGTAGTTTTCGTCGTTTGCGACTA)^9^, 500 μCi ^35^S Met-Cys (EasyTag-Perkin Elmer) and incubated at 37 °C for 90 min. All samples were treated with Benzonase (Sigma) on ice for 30 minutes to hydrolyse any polypeptidyl-tRNAs. An equal volume of gel loading buffer (186 mM Tris-Cl pH6.7, 15% glycerol, 2% SDS, 0.5 mg/ml bromophenol blue) was added to the samples and incubated at room temperature for 60 minutes. Translation products were resolved on 12% NuPAGE Bis-Tris gels (Thermo Fisher) in MOPS running buffer (50 mM MOPS, 50 mM Tris, 1 mM EDTA and 0.1 % SDS). Gels were dried under vacuum then exposed with a phosphorimager screen and scanned with Typhoon 9400 (GE Healthcare) for quantification.

## Acknowledgements

Research was supported by funding to BJB from the Academy of Finland (307431 and 314706) and the Sigrid Juselius Foundation Senior Investigator Award.

## Conflict of Interest Statement

The authors declare no conflict of interest.

## Notes

### Competing Interest Statement

The authors have declared no competing interest.

## References

1. Anderson, S., Bankier, A. T., Barrell, B. G., et al. (1981) Sequence and organization of the human mitochondrial genome. Nature, 290, 457–465.

2. Suomalainen, A. and Battersby, B. J. (2018) Mitochondrial diseases: the contribution of organelle stress responses to pathology. Nat. Rev. Mol. Cell Biol., 19, 77–92.

3. Ott, M., Amunts, A. and Brown, A. (2016) Organization and Regulation of Mitochondrial Protein Synthesis. Annu. Rev. Biochem., 85, 77–101.

4. Signes, A. and Fernandez-Vizarra, E. (2018) Assembly of mammalian oxidative phosphorylation complexes I-V and supercomplexes. Essays Biochem., 62, 255–270.

5. Kummer, E. and Ban, N. (2021) Mechanisms and regulation of protein synthesis in mitochondria. Nat. Rev. Mol. Cell Biol., 22, 307–325.

6. Vazquez-Calvo, C., Suhm, T., Büttner, S., et al. (2020) The basic machineries for mitochondrial protein quality control. Mitochondrion, 50, 121–131.

7. He, S. and Fox, T. D. (1997) Membrane translocation of mitochondrially coded Cox2p: distinct requirements for export of N and C termini and dependence on the conserved protein Oxa1p. Mol. Biol. Cell, 8, 1449–1460.

8. Hell, K., Neupert, W. and Stuart, R. A. (2001) Oxa1p acts as a general membrane insertion machinery for proteins encoded by mitochondrial DNA. EMBO J., 20, 1281–1288.

9. McDowell, M. A., Heimes, M. and Sinning, I. (2021) Structural and molecular mechanisms for membrane protein biogenesis by the Oxa1 superfamily. Nat. Struct. Mol. Biol., 28, 234–239.

10. Kumazaki, K., Chiba, S., Takemoto, M., et al. (2014) Structural basis of Sec-independent membrane protein insertion by YidC. Nature, 509, 516–520.

11. Thompson, K., Mai, N., Oláhová, M., et al. (2018) OXA1L mutations cause mitochondrial encephalopathy and a combined oxidative phosphorylation defect. EMBO Mol. Med., 10.

12. Stiburek, L., Fornuskova, D., Wenchich, L., et al. (2007) Knockdown of human Oxa1l impairs the biogenesis of F1Fo-ATP synthase and NADH:ubiquinone oxidoreductase. J. Mol. Biol., 374, 506–516.

13. Hell, K., Herrmann, J. M., Pratje, E., et al. (1998) Oxa1p, an essential component of the N-tail protein export machinery in mitochondria. Proc. Natl. Acad. Sci. U. S. A., 95, 2250–2255.

14. Hildenbeutel, M., Theis, M., Geier, M., et al. (2012) The membrane insertase Oxa1 is required for efficient import of carrier proteins into mitochondria. J. Mol. Biol., 423, 590–599.

15. Szyrach, G., Ott, M., Bonnefoy, N., et al. (2003) Ribosome binding to the Oxa1 complex facilitates co-translational protein insertion in mitochondria. EMBO J., 22, 6448–6457.

16. Haque, M. E., Elmore, K. B., Tripathy, A., et al. (2010) Properties of the C-terminal Tail of Human Mitochondrial Inner Membrane Protein Oxa1L and Its Interactions with Mammalian Mitochondrial Ribosomes. J. Biol. Chem., 285, 28353–28362.

17. Itoh, Y., Andréll, J., Choi, A., et al. (2021) Mechanism of membrane-tethered mitochondrial protein synthesis. Science, 371, 846–849.

18. Hornig-Do, H.-T., Tatsuta, T., Buckermann, A., et al. (2012) Nonsense mutations in the COX1 subunit impair the stability of respiratory chain complexes rather than their assembly. EMBO J., 31, 1293–1307.

19. Zurita Rendón, O. and Shoubridge, E. A. (2012) Early complex I assembly defects result in rapid turnover of the ND1 subunit. Hum. Mol. Genet., 21, 3815–3824.

20. Richter, U., Lahtinen, T., Marttinen, P., et al. (2015) Quality control of mitochondrial protein synthesis is required for membrane integrity and cell fitness. J. Cell Biol., 211, 373–389.

21. Richter, U., Ng, K. Y., Suomi, F., et al. (2019) Mitochondrial stress response triggered by defects in protein synthesis quality control. Life Sci. Alliance, 2, e201800219.

22. Casari, G., De Fusco, M., Ciarmatori, S., et al. (1998) Spastic paraplegia and OXPHOS impairment caused by mutations in paraplegin, a nuclear-encoded mitochondrial metalloprotease. Cell, 93, 973–983.

23. Di Bella, D., Lazzaro, F., Brusco, A., et al. (2010) Mutations in the mitochondrial protease gene AFG3L2 cause dominant hereditary ataxia SCA28. Nat. Genet., 42, 313–321.

24. Pierson, T. M., Adams, D., Bonn, F., et al. (2011) Whole-exome sequencing identifies homozygous AFG3L2 mutations in a spastic ataxia-neuropathy syndrome linked to mitochondrial m-AAA proteases. PLoS Genet., 7, e1002325.

25. Gorman, G. S., Pfeffer, G., Griffin, H., et al. (2015) Clonal expansion of secondary mitochondrial DNA deletions associated with spinocerebellar ataxia type 28. JAMA Neurol., 72, 106–111.

26. Cagnoli, C., Stevanin, G., Brussino, A., et al. (2010) Missense mutations in the AFG3L2 proteolytic domain account for ∼1.5% of European autosomal dominant cerebellar ataxias. Hum. Mutat., 31, 1117–1124.

27. Mancini, C., Orsi, L., Guo, Y., et al. (2015) An atypical form of AOA2 with myoclonus associated with mutations in SETX and AFG3L2. BMC Med. Genet., 16, 16.

28. Puchades, C., Ding, B., Song, A., et al. (2019) Unique Structural Features of the Mitochondrial AAA+ Protease AFG3L2 Reveal the Molecular Basis for Activity in Health and Disease. Mol. Cell, 75, 1073-1085.e6.

29. Battersby, B. J., Richter, U. and Safronov, O. (2019) Mitochondrial Nascent Chain Quality Control Determines Organelle Form and Function. ACS Chem. Biol., 14, 2396–2405.

30. Rep, M., Nooy, J., Guélin, E., et al. (1996) Three genes for mitochondrial proteins suppress null-mutations in both Afg3 and Rca1 when over-expressed. Curr. Genet., 30, 206–211.

31. Arlt, H., Tauer, R., Feldmann, H., et al. (1996) The YTA10-12 complex, an AAA protease with chaperone-like activity in the inner membrane of mitochondria. Cell, 85, 875–885.

32. Antonicka, H., Sasarman, F., Nishimura, T., et al. (2013) The Mitochondrial RNA-Binding Protein GRSF1 Localizes to RNA Granules and Is Required for Posttranscriptional Mitochondrial Gene Expression. Cell Metab., 17, 386–398.

33. Zhou, A., Rohou, A., Schep, D. G., et al. (2015) Structure and conformational states of the bovine mitochondrial ATP synthase by cryo-EM. Elife, 4, e10180.

34. Tsukihara, T., Aoyama, H., Yamashita, E., et al. (1996) The whole structure of the 13-subunit oxidized cytochrome c oxidase at 2.8 A. Science, 272, 1136–1144.

35. Seneca, S., Abramowicz, M., Lissens, W., et al. (1996) A mitochondrial DNA microdeletion in a newborn girl with transient lactic acidosis. J. Inherit. Metab. Dis., 19, 115–118.

36. Jackson, C. B., Huemer, M., Bolognini, R., et al. (2019) A variant in MRPS14 (uS14m) causes perinatal hypertrophic cardiomyopathy with neonatal lactic acidosis, growth retardation, dysmorphic features and neurological involvement. Hum. Mol. Genet., 28, 639–649.

37. Sanchez, M. I. G. L., Mercer, T. R., Davies, S. M. K., et al. (2011) RNA processing in human mitochondria. Cell Cycle, 10, 2904–2916.

38. Temperley, R. J., Seneca, S. H., Tonska, K., et al. (2003) Investigation of a pathogenic mtDNA microdeletion reveals a translation-dependent deadenylation decay pathway in human mitochondria. Hum. Mol. Genet., 12, 2341–2348.

39. Hejzlarová, K., Kaplanová, V., Nůsková, H., et al. (2015) Alteration of structure and function of ATP synthase and cytochrome c oxidase by lack of Fo-a and Cox3 subunits caused by mitochondrial DNA 9205delTA mutation. Biochem. J., 466, 601–611.

40. Leonhard, K., Guiard, B., Pellecchia, G., et al. (2000) Membrane protein degradation by AAA proteases in mitochondria: extraction of substrates from either membrane surface. Mol. Cell, 5, 629–638.

41. Kummer, E., Leibundgut, M., Rackham, O., et al. (2018) Unique features of mammalian mitochondrial translation initiation revealed by cryo-EM. Nature, 560, 263–267.

42. Antonicka, H., Lin, Z.-Y., Janer, A., et al. (2020) A High-Density Human Mitochondrial Proximity Interaction Network. Cell Metab., 32, 479–497.

43. Singh, A. P., Salvatori, R., Aftab, W., et al. (2020) Molecular Connectivity of Mitochondrial Gene Expression and OXPHOS Biogenesis. Mol. Cell, 79, 1051–1065.

44. Guo, M., Xu, Y. and Gruebele, M. (2012) Temperature dependence of protein folding kinetics in living cells. Proc. Natl. Acad. Sci. U. S. A., 109, 17863–17867.

45. Nakase, H., Moraes, C. T., Rizzuto, R., et al. (1990) Transcription and translation of deleted mitochondrial genomes in Kearns-Sayre syndrome: implications for pathogenesis. Am. J. Hum. Genet., 46, 418–427.

46. Parikh, S., Goldstein, A., Karaa, A., et al. (2017) Patient care standards for primary mitochondrial disease: a consensus statement from the Mitochondrial Medicine Society. Genet. Med., 19, 1380.

47. Antonicka, H. and Shoubridge, E. A. (2015) Mitochondrial RNA Granules Are Centers for Posttranscriptional RNA Processing and Ribosome Biogenesis. CellReports, 10, 920–932.

